# The reward system plays a role in natural story comprehension

**DOI:** 10.1101/2023.07.13.548681

**Authors:** Oren Kobo, Yaara Yeshurun, Tom Schonberg

**Affiliations:** Tel Aviv University

## Abstract

Prediction is a fundamental process that spans most facets of human cognition and is one of the most essential aspects of efficient language processing. At the same time, prediction plays a critical role in reward processing. Nevertheless, the exploration of the involvement of the reward system during language processing has not yet been directly tested. Here, we investigated the role of reward-processing regions while listening to a natural story. In a pre-registered study, we utilized a published dataset in which half of the participants listened to a natural story and the others listened to a scrambled version of it. We compared the functional MRI signals in the reward system between these conditions and discovered a unique pattern that differentiates between them. This suggests that the reward system is activated during the comprehension of natural stories. We also found that the fMRI signals in reward areas are related to the predictability level of processed sentences and that the system might be involved in higher predictability during the processing of a natural story.

## Introduction

The reward system has been shown to be involved in a wide variety of processes in the brain (Smillie & Wacker 2014). Prediction is a hallmark of the reward system (Schultz et al 1997). Accordingly, several theories propose that the brain’s primary objective is to reduce surprise given sensory input (Friston 2010, Den Ouden et al., 2012). Reward prediction error signals encodes the difference between actual received and predicted rewards using the phasic activity of dopamine neurons (Schultz, 2016).

In psycholinguistics, prediction is a key explanation of the human ability to comprehend language efficiently (Crystal & House, 1990; Liberman et al, 1967). The negative correlation between N400 evoked response potential (ERP) component and word predictability provides neural evidence of the relationship between predictability and speed of processing and exemplifies the major role of statistics in prediction (Kuperberg & Jaeger, 2016; Levy, 2008) and in language comprehension (Conway et al 2010; Saffran, 2003). Several fMRI studies focused on brain activity evoked by a surprising ending or semantic plausibility (Lau et al, 2008) and showed that the left inferior frontal cortex is consistently activated in such cases. Other studied unexpected input and how it influenced evoked neural activity, e.g. the effect of sentence type (Stringaris, 2005); word knowledge (Hagoort et al., 2004) or type of anomaly (Kuperberg et al., 2008; Newman et al., 2001). It was also demonstrated that sentence comprehension specifically is attributed to the middle temporal gyrus (Brennan et al 2012, Hickok 2007). However, all those studies concentrated on traditional linguistic areas in the brain and on mapping different aspects of the input and their influence on evoked neuronal activity. Here, unlike previous studies we aimed to directly test the involvement of the reward system in language comprehension of natural stories and test the effect of expectations during processing of natural stories.

Scrambling of sentences is a tool often utilized to distill cognitive semantic processes when contrasted with corresponding intact sentences. It was used to explore properties of timescale hierarchy and temporal accumulation of information during processing across the cortex. It has been shown that the longest timescales were for default-mode networks, shorter for intermediate areas along the superior temporal gyrus and the shortest to early sensory regions (Hasson et al 2015; Zuo et al 2020; Lerner et al., 2011; Yeshurun et al 2017). Recently, Natural Language Processing (NLP) computational modeling has become more commonly used in the investigation of processing during comprehension of a naturalistic story. Several studies correlated a predictability-related metric with various facets of neural activity while listening to natural stories (Frank & Willems 2017, Willems et al 2015). Using stories rather than controlled stimuli has many advantages. For example, they can be used to explore various questions in a single dataset, and often reveal more widespread responses to language than controlled stimuli (Tikochinski et al 2023; Hamilton & Huth 2018; De heer et al 2017; Huth et al 2016). Computational tools are now used to formalize the attributes of the process in question and correlate it with activity to gain a better understanding of underlying cognitive processes (Willems et al 2015 used surprise and entropy; Frank & Willems 2017 semantic distance, and Lopopolo et al 2017 perplexity).

Therefore, here we used sentence perplexity to test the role of the reward system during comprehension of sentences. We analyze an openly shared fMRI dataset where participants listened to a story and a scrambled version of the same story. We surmise that given the prominent role of the reward system in generating and calibrating predictions, and the role predictions play in language processing, the reward system is likely to take part in sentence comprehension. We hypothesized we will be able to distinguish between conditions based on activity in the reward system and exemplify that the nature of this distinction can be attributed to sentence predictability. Our findings will test if the predictability of the linguistic input elicits processing in the reward system, which to date has not been directly demonstrated.

## Materials and Methods

This study was pre-registered in: https://osf.io/mt7yd/. Analyses outside of the pre-registration are mentioned specifically in the text. We performed all analyses within a pre-registered reward mask and a control area (within the visual network), both obtained from Neurosynth.org (Yarkoni et al. 2011) by entering the corresponding terms in the meta-analyses tool and downloading the association map. We used classification and computational models to probe whether there was a different response in the reward network to the experimental conditions of a natural story (intact) and a scrambled version of it (scrambled). If indeed this was the case, we aimed to inspect the relatedness of these findings specifically to predictability.

All analysis code are published: https://github.com/orenpapers/Reward_Predictability_Paper

### Stories and experimental design

The current study reanalyzes a previously published fMRI dataset (Yeshurun et al., 2017) that was shared as part of the narrative dataset (Nastase et al, 2020). In the dataset reanalyzed in this study, 36 right-handed participants listened inside the MRI scanner to one of two narratives: 18 participants listened to the *intact* story and ;18 other participants listened to the *scrambled* story. The intact story was about a woman that was obsessed with an American Idol judge, meets a psychic, and then becomes fixated on Vodka. The *scrambled* story had the exact same words as the intact, but the words within each sentence were randomly scrambled (Fig. 1). This manipulation created an unpredictable input throughout the *scrambled* story. Both stories were read and recorded by the same actor. The beginning of each sentence was aligned post-recording. Each story was 6:44 minutes long and was preceded by 18 seconds of neutral music and 3 seconds of silence. Stories were followed by an additional 15 seconds of silence. These music and silence periods were discarded from all analyses (see the Analysis section below). Experimental procedures were approved by the Princeton University Committee on Activities Involving Human Participants. All participants provided written informed consent.

**Figure 1:**
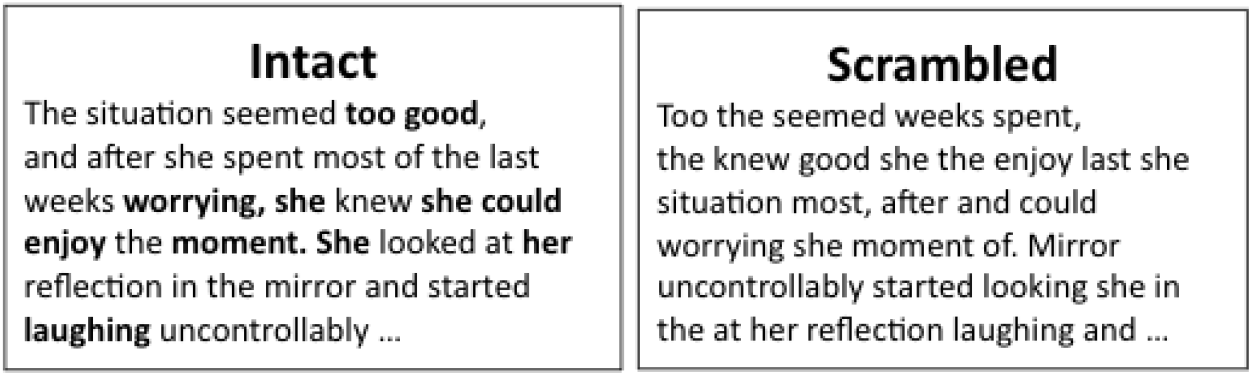
Edited from (Yeshurun et al, 2017). We used 2 of the 4 conditions of the original study. Intact story had the same words as Scrambled, but the words were scrambled within a sentence. Thus, there was no words change, but scrambled words that resulted in unpredictable input

### Sentence predictability

The effect of prediction on comprehension and generation of expectation has been shown in the syntactic (R. Levy, 2008) and phonetic (Marslen-Wilson, 1975) expectations. In the current study, we chose to focus on semantic predictability (the probability of a certain word in context regardless of the structural structure). This approach was used in various cases to formalize predictability by statistical expectation, for example by using statistical measures such as transitional probability and the likelihood of two words to co-occur. We measure sequence predictability using perplexity. The perplexity (PP) of a language model on a test set is the inverse probability of the test set, normalized by the number of words (See eq. 1). Thus, minimizing perplexity is equivalent to maximizing the test set probability according to the language model. (Jurafsky & Martin 2009)

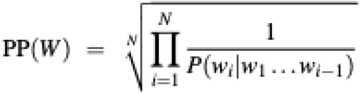

### Statistical analyses

In all the analyses, we calculated an accuracy level for the intact condition by running cross-validation folds. Thus, we generated a global accuracy score and a distribution of accuracy per fold. In addition, we also compared actual accuracies to a corresponding controlled null distribution, generated by either using the data from the scrambled condition or by shuffling the actual labels. To test whether a result was significant, we compared the actual result with the null distribution results. The p-value using the rank of the mean accuracy of the actual data folds in the null distribution. Then, we ran the exact same analysis on the preregistered control area (visual-related region). We set the significance level in our analyses to p=0.05 and added FDR multiple comparisons correction when relevant. All statistical tests were performed in Python.

### Differentiating between activity in intact vs. scrambled conditions

First, to test whether the reward system is involved in language processing, we tested if there was a difference in its brain response between the intact and scrambled conditions. For that, we applied the following analyses:

#### 1. Condition classifier at the participant level

In this pre-registered analysis, we trained a support vector machine model (SVM) classifier at the participant level to predict a condition given participants’ activity. For that purpose, a participant activity vector (sized 1X66) was created by averaging responses across sentences, such that this vector provides a representation of the neural activity of the participant during the story (See Figure 2A). We then ran 1000 iterations with 5-fold cross-validation. In each iteration, we trained the model on the training participants and evaluated it on the test data participants, as assigned by the cross-validation. Due to the high dimensionality of the data relative to the number of samples, we first applied a PCA with 20 components to the data. Then, we obtained p-values using a permutation test (artificially generated by running 1000 iterations of the model with shuffled labels). We applied the exact same procedure for the pre-determined control vision area.

**Figure 2:**
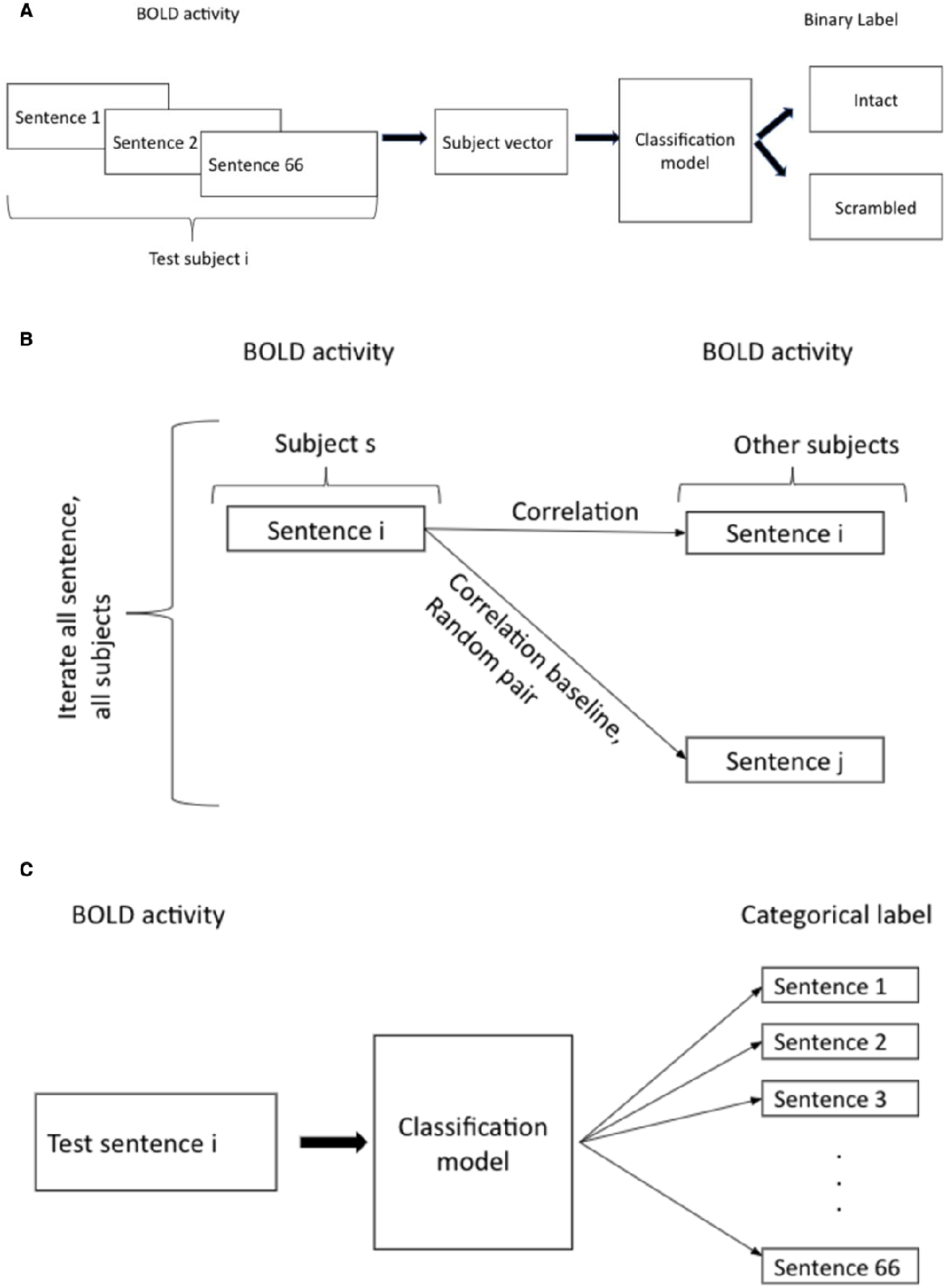
**A**: The process of creating a classifier for each participants’ condition. We averaged neural activity, per sentence, throughout the task, to generate a single vector per participants with the width of 66. This was used as the input for the model, and the output for a binary label that was the condition. **B**: Between-participants pattern similarity of sentences. For each sentence, neural data was averaged across time within each voxel, resulting in one vector for each sentence. Correlations were computed between sentences across participants. The null distribution was generated by choosing a random sentence, in addition to the actual pair. **C**: multiclass sentence classifier. In each fold, we trained the model to predict one of the 66 experimental sentences based on elicited neural activity. We then predicted the correct labels for each of the 66 sentences of the test participant.

#### 2. Euclidean Distance Measure

In this pre-registered analysis, we aimed to evaluate the extent to which activity in reward areas alternates between conditions. To do so, we used the Euclidean Distance Measure which provides a measure (per voxel\ROI) of the difference in the neural response between two conditions (Yeshurun et al, 2017). We measured the significance of the difference between the activity of the 2 conditions, per voxel, as follows: For each condition, we created a matrix of mean activity (across participants) per TR, per voxel – which resulted in 2 matrices with the shape of n_TR X n_voxels. Then, for each voxel, we calculated the Euclidean distance between its respective time series -to get a distance vector with the shape of n_Voxels. Thus, at each point, the vector stores the Euclidean distance (over time) of the averaged (across participants) distance. Afterward, we ran 1000 iterations of label shuffling, to obtain the null distribution of 1000 artificial distances vectors. Finally, for each voxel, we calculated the rank of the real value compared to the null distribution and calculated z-scores based on this rank and adjusted the p-value for each voxel using False Discovery rate method (Benjamini & Hochberg 1995). Moreover, we compared the percentage of significant voxels in the reward system and the control vision area.

### Sentence-Level information in the reward system

we tested whether there was any information regarding the sentence themselves that was encoded in the reward system. Such a finding would support the suggestion that the reward system was involved in the processing of stories by showing its activity was directly related to actual perceived language information. We ran the following analyses:

#### 1. Pattern similarity between sentences

We tested if the same sentence elicited similar activity across all participants in the reward system by using an analysis that measures between-participants spatial similarity (Chen et al 2016). We iterated over all sentences and all participants. In each iteration, we calculated the correlation between the activity from the specific pairings of participants and the sentence to the activity elicited by the same sentence in all other participants. Additionally, to create a baseline null distribution, an additional sentence was randomly selected in every iteration and its correlation with the original sentence was calculated (See Figure 2B).

#### 2. Multisentences classifier

we measured the extent of sentence-level information encoded in the reward system during processing by probing if we could train a classifier to predict the sentences currently processed. We ran iterations of Leave-One-Out participants in the intact condition and trained a multilabel classifier (66 labels) on 17 train set participants. We then used the trained model to classify the 66-width vector (each entry was neural activity elicited by a specific sentence) of the test participants and obtained an accuracy score (See Figure 2C). For each participant, there were 66 sentences and 66 corresponding labels and therefore this was a multiclass classification. Similarly to the condition classifier, we applied PCA with 20 components followed by an SVM and generated a null distribution with 1000 iterations of randomized labels, in order to obtain the significance level.

### Probing the relation between activity and predictability

We further conducted the following analyses to accommodate the hypothesis that activity in the reward system during processing was related specifically to the predictability of the linguistic input, and not to other possible aspects (e.g. narrative content) that differs between condition. Therefore, these analyses were designed to characterize specifically the relation between reward activity and sentence predictability to better establish the claim that differentiating activity between the conditions is associated specifically with predictability. In that regard, we performed the following analyses:

#### 1. Ordinal Classification of sentence perplexity

We used ordinal regression to predict the rank of sentence predictability within the sentences vector. We chose the approach of ordinal regression as this variable has an arbitrary scale where only the relative ordering between different values is significant. Similarly to the Multisentences classifier described above, here we also ran iterations of Leave-One-Out participants in the intact condition. We trained a multilabel classifier (66 labels) on 17 train-set participants and tested the prediction on the remaining test participant. We trained the classifier to estimate the perplexity of a given sentence based on the neural activity it elicited. We averaged per-sentence activity across participants to yield a single vector per sentence and again applied a PCA. We used mean squared error (MSE) as a metric to evaluate performance and compared the MSE of the model trained to a model trained on the scrambled version of the story. Meaning, to obtain the p-value we ran this procedure 1000 times on the scrambled data, to generate the null distribution for the permutation test. This was done to ensure that any possible effect is to be attributed to predictability rather than words themselves. To assess statistical significance, we used results obtained from the scrambled condition as the null distribution of the classifier.

#### 2. Across participant perplexity classifier

In this pre-registered analysis, similarly to the previous one, we aimed to separate plausibility from predictability and assessed the extent that reward activity was related to sentence predictability. We trained a classifier to distinguish between above and below-median sentences within the intact condition, across all participants. Thus, for each sentence in the intact condition, we stacked all the neural response signals from all the participants together. We then divided the sentences into low and high predictability (above/below median) using perplexity scores yielded from the BERT Language model. Finally, we trained and evaluated 1000 iterations of train folds (20% test set) and obtained p-values by comparing to the null distribution. This was done separately for reward and visual areas.

#### 3. Within participant perplexity classifier

In this pre-registered analysis, we aimed to test individual differences in the relation between reward areas and predictability. We divided the sentences to low and high predictability (above/below median) using perplexity scores yielded from the BERT Language model. Then, for each participant’s data, we trained and evaluated 1000 iterations with a 20% test set and obtained p-values by comparing them to the null distribution. This was done separately for reward and visual areas.

### Whole-brain searchlight analysis

We also applied an exploratory analysis, in which we applied the Euclidean Distance measure procedure at the whole-brain level (and not within the specific pre-registered ROIs). For each voxel, we evaluated the extent to which it was indicative of one of the two conditions by assessing the z-score of the actual distance between neural data between conditions and the null distribution. To establish the cognitive functions in which these regions were most involved, we conducted a formal reverse inference analysis using NeuroSynth (Yarkoni et al 2011), correlating our searchlight map with the neural activation maps for each term in the NeuroSynth database.

## Results

### Differentiating between activity in intact vs. scrambled conditions

We used an SVM classifier to assess differentiation between activity in the response to the intact and scrambled stories. Based on the response of the reward system, there was an accuracy of 74% in predicting the condition. This accuracy value was significant according to the permutation test (p=0.012, see Figure 3A). As a control, we also ran the same analysis based on activity in the visual system, for which we received an insignificant result in a permutation test (p = 0.085). Although a p-value of .085 is approaching significance, note that no voxel in the visual system survived the following analysis we conducted to measure differentiation between the conditions, unlike in the reward system.

**Figure 3:**
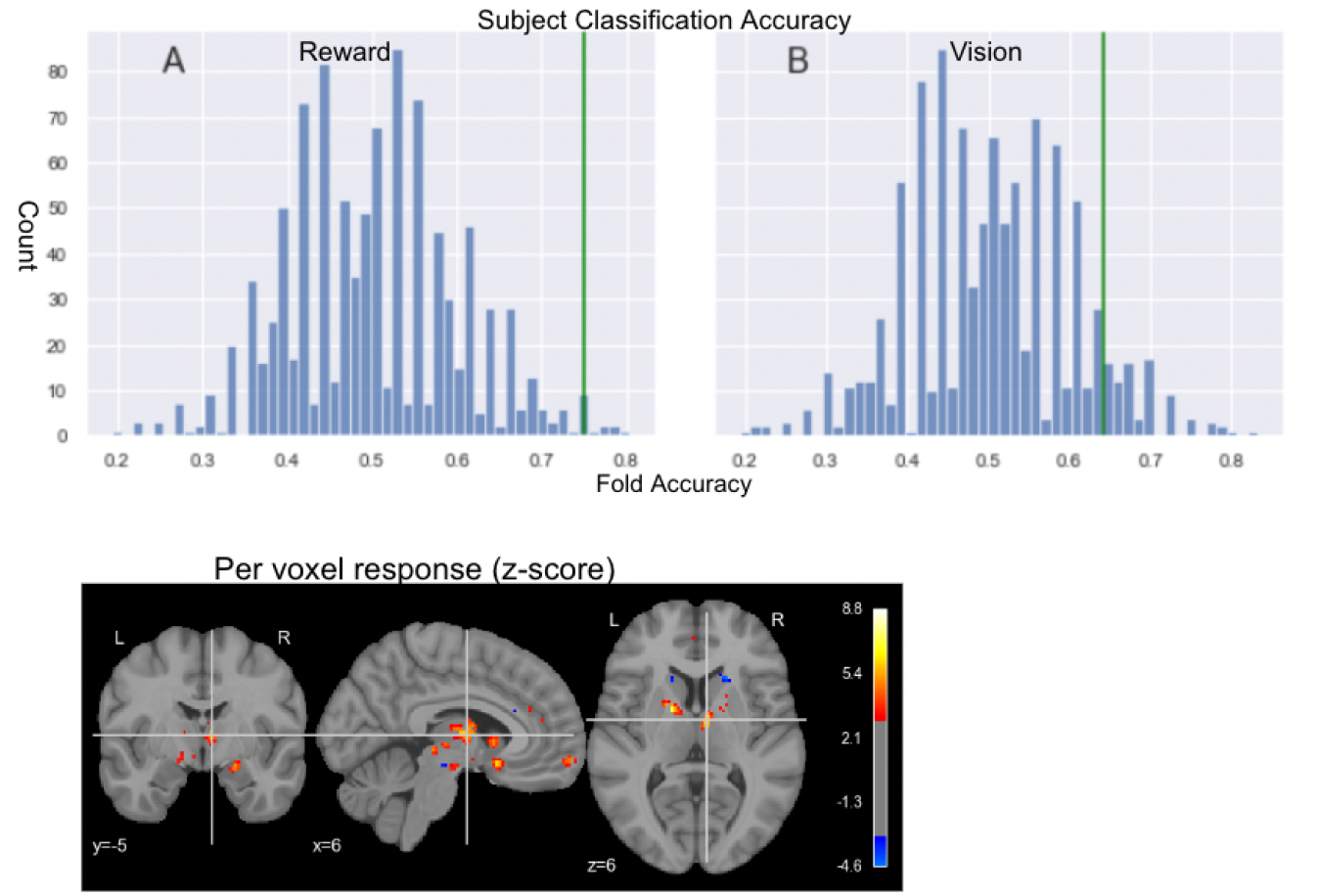
UP: Results of condition classification per participant: rank of accuracy score in the null distribution results (permutation test). The null distribution was generated by shuffling of the labels, to obtain p-values. A: reward system B: vision system (control). Down: per-voxel difference (z score) between neural response to the intact vs. the scrambled story at (x=6.44, y=-5.63, z=6.71)

To further explore the differentiation between conditions, we calculated the Euclidean distance between the averaged time courses for the intact and scrambled conditions in each voxel across the reward system (12031 voxels). For each voxel, we obtained the z-score of the intact-vs-scrambled Euclidean distance, compared to the null distribution, and calculated the adjusted p (with FDR). This analysis revealed 268 voxels in the reward system in which there was a significant difference in the response to the intact vs. the scrambled story (See Figure 3B) In the control visual system, this analysis revealed no such voxels.

### Encoding of sentence-level information in the reward system

We conducted pattern similarity analysis – i.e. we correlated activity between sentences in the reward mask to detect an intra-participant correlation of evoked activity per sentence in the reward system. This was done to test for shared aspects of processing of sentence across participants in the reward system. We obtained a weak but significant correlation of 0.09 for the actual data compared to the 0.01 for the baseline (randomly chosen pairs) (see Figure 6). We did not find a significant correlation between perplexity and pattern-similarity per sentence. Meaning, there was no relation between values pattern similarity of sentences to perplexity values. We also tested pattern-similarity per-participant and per-sentence to assess the robustness of such a correlation. At the participant level, we found that for each of the participants there is a significant difference between the spatial correlation of elicited activity in the real compared to randomly chosen (baseline) pairs (p=0.04 for a single participant and p < 0.01 for all others). At the sentence level, we found a significant correlation only for 34 out of the 66 sentences. This may imply that activity in the reward system is moderated or activated by certain sentences (or types of sentences) to a higher extent than by others. Guided by this finding, we also trained a sentences classifier. The classifier was trained to predict the processed sentences based on the activity in the reward area. We obtained an accuracy of 0.053 (while chance level is 0.01515 since there are 66 sentences). This was significantly above chance level (p = 0.016) in the permutation test, whereas for the control visual network and control scrambled condition we obtained non-significant results (p = 0.28 and p=0.29 respectively) (see Fig 4). We did not find correlations between sentence perplexity and classification accuracy. Meaning, level of perplexity did not influence performance of the model. However, there was a strong correlation between accuracy of sentence classification and the pattern similarity of a sentence – (spearman r = 0.35, p = 0.003), meaning that sentences that got higher accuracy rates also tended to have higher pattern similarity. This could further indicate that certain sentences eliciting activity in reward regions in a different manner than other sentences.

**Figure 4:**
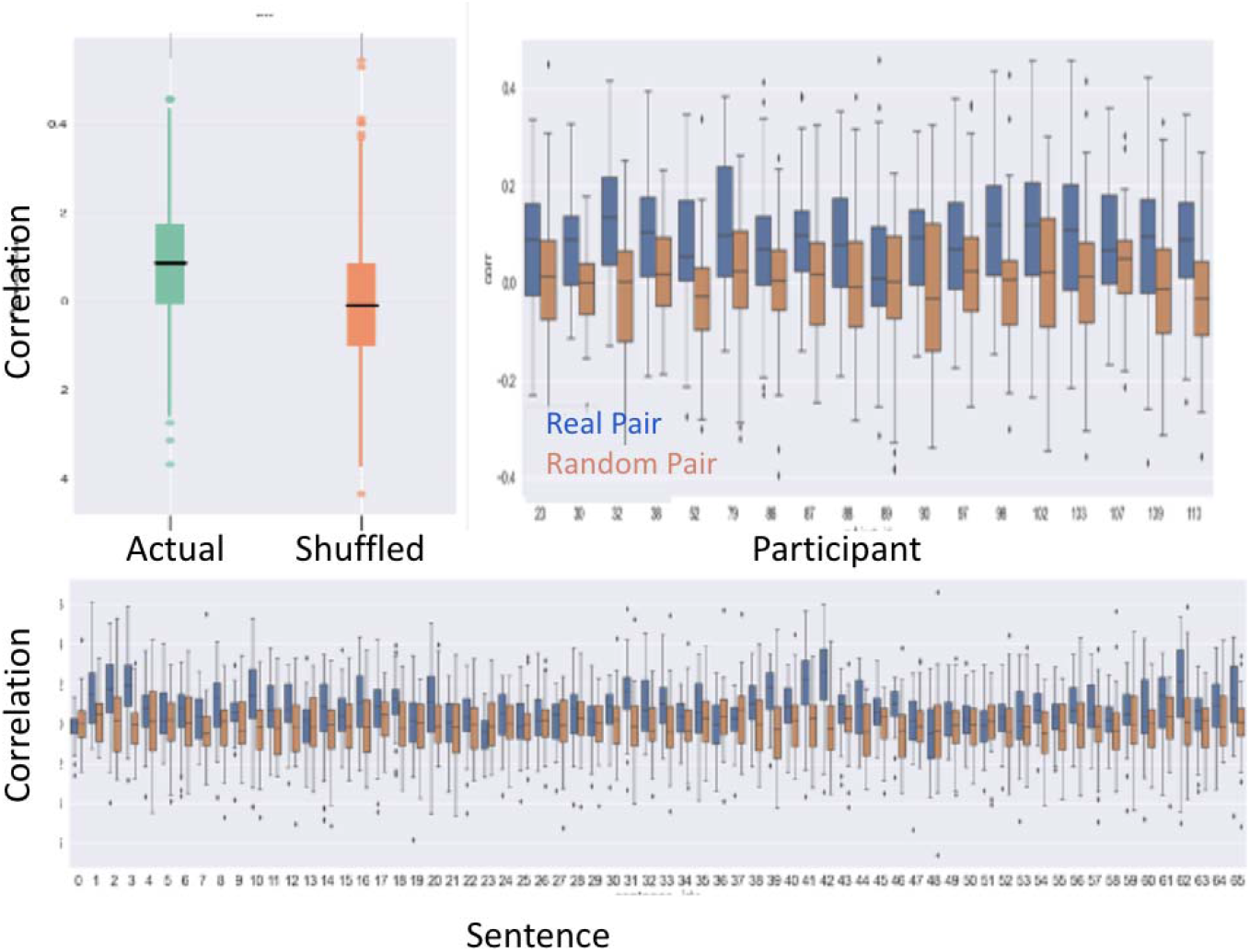
A: Correlation of elicited activity per sentence is higher for actual data compared to baseline. B: Results per participants. C: Results per Sentence.

**Figure 5:**
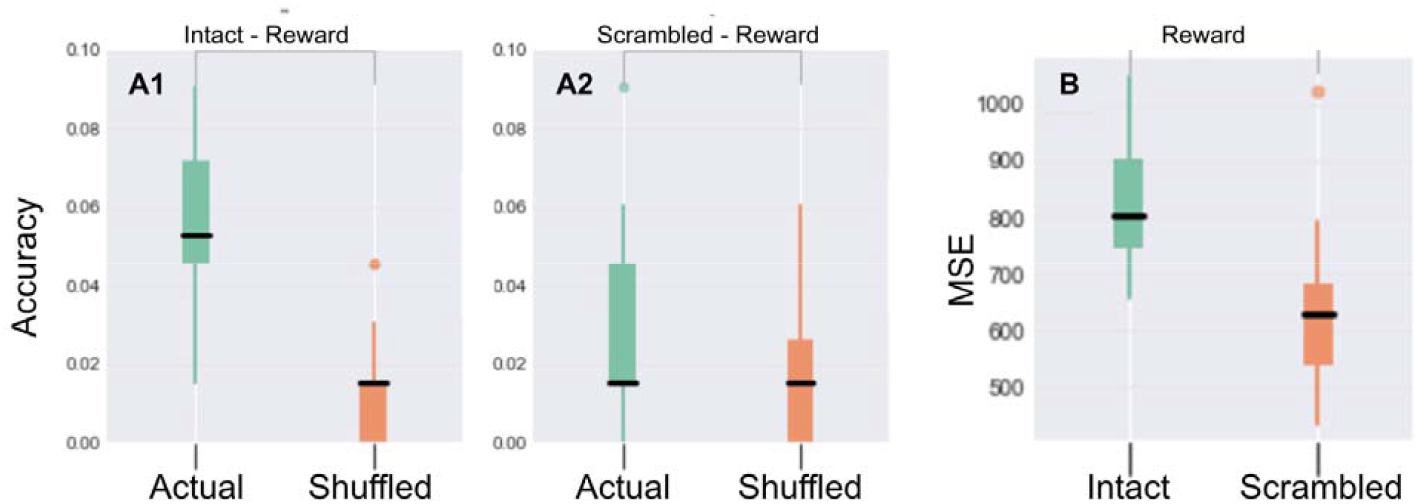
A1: Distribution of per fold accuracy (actual vs permutations) for the intact condition. A2: Distribution of per fold accuracy (actual vs permutations) for the scrambled condition. B: Distribution of per fold MSE of the intact condition, compared to scrambled condition for the ordinal classification. All models are based on neural data acquired in the reward system.

**Figure 6:**
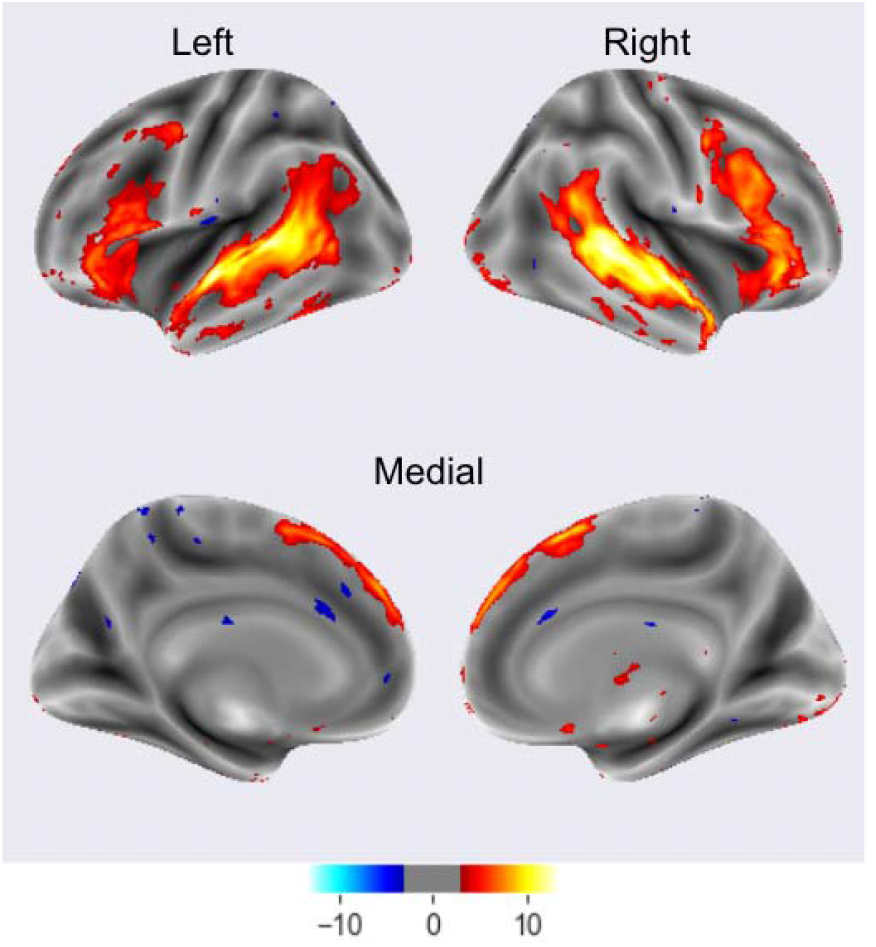
Voxels activity difference between scrambled and milky conditions (z score)

### The relation between activity and predictability

We examined if the relation between activity in the reward system could be attributed to predictability. Meaning, to probe whether this activity could be attributed to differences in predictability between conditions, or to an alternative explanation (e.g. the existence of a structured narrative in the intact condition or the rewarding nature of the words themselves).

#### Ordinal classifier for sentence perplexity

We trained an ordinal classifier for sentence perplexity, based on activity in the reward system in the intact condition and in the scrambled condition as control and obtained an MSE = 625 (p < 0.0001) in the permutation test for the reward system, whereas in the control condition, results were insignificant (p=0.44). (See Fig.5). *Across participants perplexity classifier:* In this pre-registered analysis we trained a binary classifier to predict the perplexity of a sentence (above or below the median) given elicited neural data from the reward system in the intact condition and compared results to the baseline of the scrambled condition. We obtained insignificant results in the permutation test, meaning that the classifier we trained to predict perplexity level of a sentence given the neural data it elicited in the reward system, did not achieve above chance level results.

#### Within participant perplexity classifier

In this pre-registered analysis we trained a separate classifier per participant, to predict the perplexity level of a given sentence, from the evoked activity while it is being processed. We did not obtain any significant result in the permutation test, meaning when trained a separate classifier per participant (instead on the aggregated data of all the participants), we also could not successfully train a classifier to predict whether a certain sentence had above median perplexity based on acquired data in the reward system.

### Whole-brain searchlight analysis

We applied a voxel-level Euclidean Distance whole brain analysis to test for regions that had a significant difference in their response to the intact and scrambled stories (Figure 6). As expected based on previous studies and the language-comprehension nature of the difference between the two stories, this exploratory analysis revealed mainly regions involved in language and comprehension processes (Table 1).

**Table 1:** Pearson correlations between searchlight classification maps and NeuroSynth term-based reverse inference activation maps obtained by uploading the map to Neurosynth. The 10 most highly correlated terms are listed.

| **Term** | **Correlation (r)** |
| --- | --- |
| Temporal sulcus | 0.444 |
| Superior temporal | 0.441 |
| Temporal | 0.429 |
| Linguistic | 0.424 |
| Sentences | 0.421 |
| Language | 0.413 |
| StS | 0.399 |
| Spoken | 0.397 |
| Comprehension | 0.30 |
| Temporal gyrus | 0.375 |

## 4. Discussion

In this study we explored the role of the reward system in language predictability during narrative comprehension. We reanalyzed a published dataset, composed of two conditions – a natural story and a scrambled version of it (Nastase et al 2021). We found that the reward system was involved in processing the intact (but not the scrambled) story. Moreover, using a sentences classifier and pattern similarity analysis we found that the reward system encoded information at the sentence level. We provide an indication that this was related to the predictability of the language input, by assessing the dynamics through which fMRI responses were affected specifically by the predictability (formalized by perplexity) of the processed sentence.

We found that the activity in the reward system was moderated by whether participants listened to an intact story, or to a scrambled version of it. The reward system has been shown to be activated during language processing, mostly in the context of a presence of an external reinforcer (Kaltwasser et al 2013; Boyer 2017; De Loof et al 2018) and humor (Shibata et al 2014; Mobbs et al 2003) or a pleasing content (Bohrn et al 2013). Semantic processing and the reward system has also been linked through the finding of spread of semantic priming activation, mainly using patients with Parkinson’s Disease that display abnormal activation (Kischka et al. 1996; Ole Tiedt et al 2022). However, it has yet to be exemplified that the reward system is activated during normal comprehension of a natural story. The ability to consistently differentiate test participants at an above-chance level between conditions based on activity in reward areas suggests that processing of the story is performed by these regions. Prediction of upcoming input is a core strategy in the brain, activated across many cognitive domains, including perception, sensory-motor processing, and learning (for reviews see Clark, 2015; Hohwy, 2018). The reward system takes part in predictions using prediction error signals (Schultz 2016), which also seem to be crucial in language comprehension due to the fast processing required (Ehrlich & Rayner, 1981). Thus, a successful prediction leads the system to improve and facilitate faster reaction times.

We further tested the hypothesis that the activation in the reward system is related to the actual processed sentence at a certain time. We found an increased pattern similarity between sentences and above chance accuracy of a classifier that predicted the processed sentence from neural activity, suggesting there was a distinct sentence-level representation. While a correlation between accuracy in sentence classification prediction to the predictability of a sentence, was not found, we did find a correlation between accuracy in sentence classification and levels of pattern similarity (meaning, sentences that tend to have higher pattern similarity tend to have higher classification accuracy). This further implies there are specific types of sentences that are better represented in the reward system. Given these findings, we conclude that there might be some extent of sentence-level information encoded in the reward system, and that the reward system encoding might be related to specific attributes of the sentence. As far as we know, this is the first time that a relation between sentence-related attributes and representation in the reward system has been exemplified.

As the two main factors that are modified between the two conditions (intact and scrambled) were the predictability of the input and the existence of a structured narrative, we further demonstrated the influence of predictability on neural activity, in order to establish that the driving factor between the alternation in elicited activity is predictability. We relied on a perplexity estimation as an indicator of sequence predictability. While it is common to use entropy and surprise, measures adapted from information theory to gauge predictability, each of them only captures a specific sense of word predictability. Perplexity is a measure taken from the field of NLP that should arguably better model sentence probability and was also utilized in cognition (Lopopolo et al., 2017). We show evidence that the neural activity in the reward system might be related to predictability by successfully training an ordinal regression classifier and a binary classifier to predict the level of perplexity given participants’ data. We provide supporting evidence that the difference in the reward system between conditions is not to be associated with changes in attention/arousal etc., in addition to activation related to comprehension per se. We exemplify it by that that the activity is related also to perplexity, and that there is also a sentence-level information in the reward system. While the per-participant classifier failed to predict the level of perplexity, it is reasonable that this was due to the low amount of data.

The human brain constantly predicts what stimulus is expected based on current information (Clark, 2013; Friston, 2005). The study of neurolinguistics and characterization of its underlying neural mechanisms identified different regions related to facets of sentence comprehension (Friederici, 2002; Hagoort, 2005) and it has been shown that prediction is a key component in it, using reading times (Goodkind & Bicknell, 2018). Functional MRI was also commonly used (Frank & Willems, 2017; Lopolopo et al 2017; Willems et al 2016), to study prediction (through semantic similarity, entropy, and surprise) under the domain of neurolinguistics. Predictability measures have been correlated with neural signals evoked during both reading and listening language comprehension tasks (Carter et al., 2019; Frank et al., 2015; Frank and Willems, 2017;). The reward system is known to take part in prediction and the prediction error signal (Shultz et al 2016). However, a link between the reward system and predictability during language comprehension has yet to be found. Here, we unveil the plausibility of such a link by showing the ability to predict levels of statistical predictability of sentences in the intact condition based on activity in reward areas.

The role of the reward system in language processing might be manifested by optimizing representation to support better prediction, leading to faster processing. An alternative possible explanation is that the role of reward is related to the existence of a narrative. This is evident by showing relatedness between activity in reward to specifically perplexity, which would not be the case if the mere existence of a narrative influences the difference in activity between the conditions. Future work may use a controlled stimuli set to better account for this alternative explanation, and explore the effect of predictability on evoked activity and how it is affected by various attributes of the processed sentence, and the extent that semantic data is represented in reward areas. Future work could also concentrate on the specific attributes of this involvement, specifically the proposed representation of sentence-related information in the reward system, aiming to define what types of sentences are represented and how, and the relation to language representation and connectivity with this area during processing.

While using natural stimuli may increase complexity, add possible artifacts or decrease effect size, it entails other advantages. Natural language studies often reveal much more widespread (and less left-lateralized) responses to language than studies with controlled stimuli (Hamilton & Huth 2018). Natural language studies also seem to be more sensitive as they could be used for exploring many different questions using a single dataset. (Hamilton & Huth 2018; De heer et al 2017; Wehbe et al. (2014) showed that different aspects of story processing were encoded in different brain networks). Therefore, although many previous studies used tightly controlled stimuli, such as a set of single isolated words, sentences or curated passages, and explicit lexical semantic tasks (Chee et al., 1999; Michael et al., 2001; Buchweitz et al., 2009; Liuzzi et al., 2017) we chose to use natural stories to provide initial evidence to our claim.

Overall, we demonstrated for the first time the involvement of the reward system during natural language comprehension, showing that the reward system plays a role in story processing. We found evidence indicating that this role is related to predictability. We propose an interpretation by which the more predictable nature of the intact condition caused this effect and that its response is related to the predictability level of the input, implying that the reward network activity might reflect a rewarding nature of language input with higher predictability. We found strong evidence for this relation in the between per-sentence explicitly predictability measure of perplexity and reward within the intact condition, but not in the scrambled condition. Future work should be done to better interpret the precise characteristics of the role the reward system plays in language processing, its specific mechanism, and underlying factor, to better characterize its relation to predictability and other attributes of the processed sentence, and specifically to dissociate it from merely narrative-related effects. Using a more controlled design can be fruitful to effectively identify the role of the reward system in comprehension

